# Circuit mechanisms underlying sexually dimorphic outcomes of early life stress

**DOI:** 10.1101/2024.11.27.625736

**Authors:** Cheng Jiang, Ignacio Ruiz-Sanchez, Cindy Mei, Christopher Pittenger

## Abstract

Stress during early life influences brain development and can affect social, motor, and emotional processes. We describe a striking sex difference in the effects of early life stress (ELS), which produces anhedonia and anxiety-like behaviors in female adolescent mice, as reported previously, but repetitive behavioral pathology and social deficits in male adolescent mice. Notably, this parallels sex differences seen in the prevalence of psychiatric symptoms: depression and anxiety disorders are more common in girls and women, whereas neurodevelopmental disorders like autism spectrum disorder and Tourette syndrome are markedly more common in boys and men. We characterized the effects of ELS on the medial prefrontal cortex (mPFC) and on its projections to the dorsal striatum (dStr) and lateral septum (LS). ELS males, but not females, developed hyperactivity in the cortico-striatal circuit and hypoactivity in the cortico-septal circuit. Chemogenetic manipulation of cortico-striatal projection neurons modulates repetitive behavioral pathology and social behaviors in stressed males, and anhedonia in stressed females. Activation of cortico-septal projection neurons rescues social deficits in stressed males. We conclude that early life stress produces sexually dimorphic behavioral effects, with potential relevance to human psychiatric symptoms, through its differential effects on cortico-striatal and cortico-septal circuits.

## INTRODUCTION

Early life physiological and psychological stress can have profound effects on brain and behavior, including the development of psychiatric symptoms later in life. This has been most intensively studied in the context of mood and anxiety dysregulation, and related phenomena.^1-3^ Notably, females are generally more susceptible to these effects of early life stress (ELS) than males;^4^ intriguingly, this parallels the observation that major depressive disorder (MDD) and most anxiety disorders show a notable female preponderance.^5^ Developmental stress can also influence neurodevelopmental disorders (NDDs), such as autism spectrum disorder (ASD) and Tourette syndrome (TS). For example, severe early life deprivation and neglect produce social deficits and repetitive behaviors.^6-14^ Stress early in development, and even during gestation, has been linked to risk of TS and to tic severity.^15-17^ Of note, ASD and TS are markedly more common in males, with a sex ratio of approximately 4:1.^18,19^ These observations suggest the possibility that developmental stress may have different effects in males and females, with potential relevance to sex differences in neuropsychiatric diseases.

The medial prefrontal cortex (mPFC) is sensitive to the effects of early life threat and deprivation;^20^ it and its subcortical projections have been implicated the pathophysiology of both MDD^21^ and NDDs.^22-25^ Neuroimaging studies have revealed dysregulated functional and structural connectivity in cortico-striatal circuits in individuals with ASD and TS,^26,27^ and this projection is involved in the development of dysregulated repetitive behavior in a number of model systems.^28-31^ Recent preclinical data also implicate the circuit between the mPFC and the lateral septum (LS) in the regulation of social behaviors.^32^ How ELS influences the development of these circuits has not been investigated.

ELS paradigms in rodents seek to model the influence of developmental stress on brain development, structure, and function. Most studies have focused on the impact of ELS on depression- and anxiety-relevant behaviors and corresponding circuitry, especially the hippocampus and amygdala, and the mPFC’s connections with them. Few studies have examined other circuits and behavioral domains, and even fewer have explored sex differences in ELS effects. Here, we characterize sexually dimorphic behavioral outcomes following ELS in mice.

## RESULTS

### ELS produces sexually dimorphic behavioral phenotypes

We applied an ELS paradigm consisting of daily maternal separation and reduced nesting materials to C57BL/6 wildtype mice.^33^ In a pilot experiment, we administered ELS during two different postnatal exposure windows, from postnatal day (PND) 2 to 10 (this group is denoted PND2S) or from PND10 to PND18 (PND10S), and tested mice in a range of behavioral assays in adolescence and adulthood **(Figure S1A**). PND2S, but not PND10S, increased grooming, impaired motor learning on the rotarod test, and impaired prepulse inhibition in adolescent male mice, but not in adolescent females or in adults (**Figure S1**). To better understand this sex difference, we focused on PND2S in subsequent experiments.

In a separate cohort of mice, PND2S ELS produced elevated grooming (**Figure 1A**) and reduced social interaction time (**Figure 1B**) and sociability (**Figure 1C**) in male but not female adolescents; these behaviors have been interpreted as capturing aspects of the pathophysiology of ASD^34^ and TS.^35^ Past work using a similar ELS paradigm has focused on its impact on depression- and anxiety-like behaviors and has found greater effects in females than in males.^4^ We therefore tested these mice in the sucrose preference test (SPT, an assay of depression-like anhedonic behavior^36^) and in the novelty suppressed feeding test (NSFT, an assay of anxiety-like behavior^37^). We observed depression- and anxiety-like behaviors in females but not in males (**Figure 1D,E**). Locomotor activity was not affected (**Figure 1F**).

**Figure 1.**
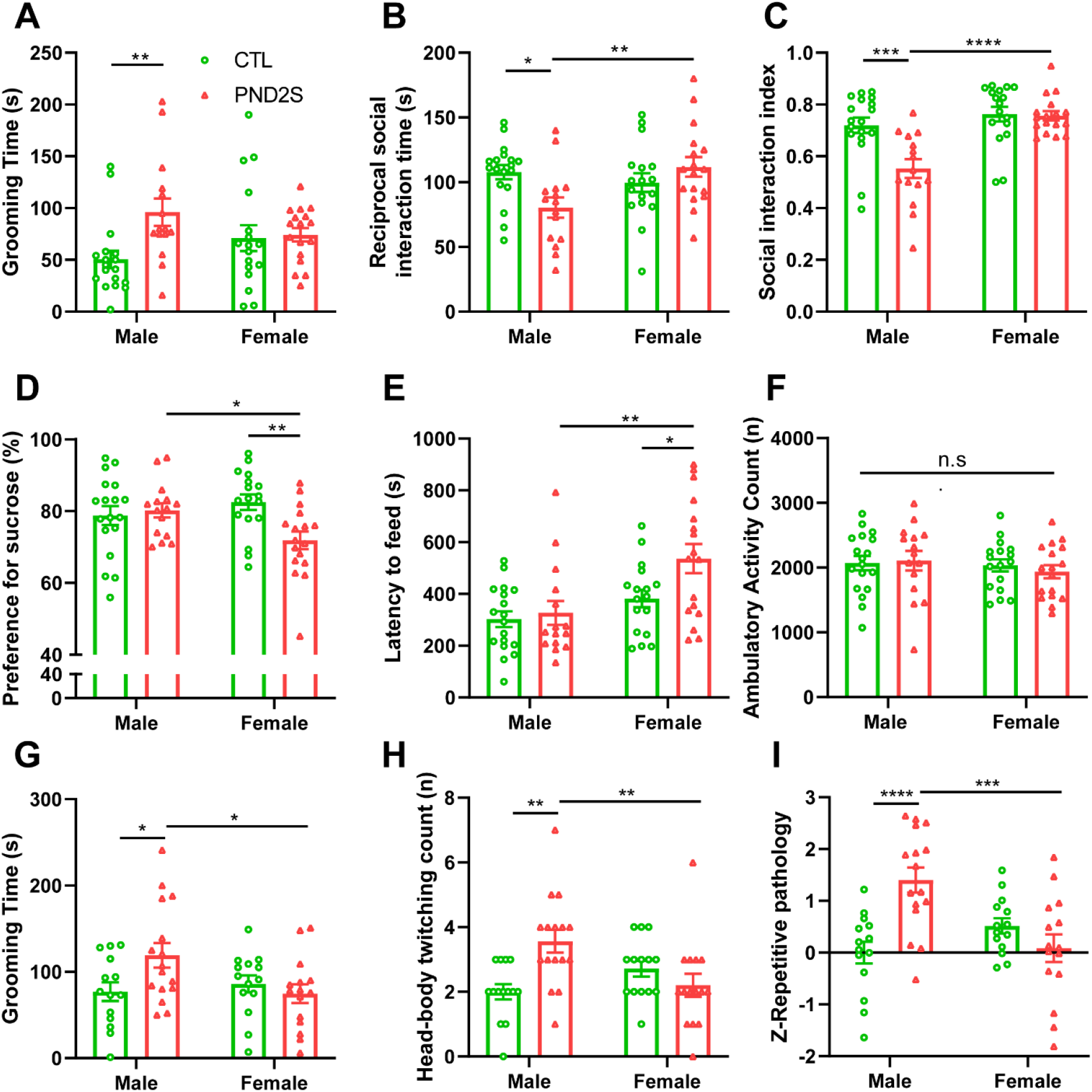
Double dissociation of the behavioral effects of ELS in males and females. **(A)** Grooming was elevated in male but not female adolescent ELS mice (Sex × ELS F[1, 63]=4.207, p=0.044). **(B)** Reciprocal social interaction was reduced by ELS in males but not females (Sex × ELS: F[1, 63]=7.9, p=0.007). **(C)** In the three-chamber sociability test, ELS reduced social preference in males but not females (Sex × ELS: F[1, 63]=8.0, p=0.006; Sex: F[1, 63]=18.9, p<0.0001; ELS: F[1, 63]=9.3, p=0.003). **(D)** In the sucrose preference test, ELS produced reduced sucrose preference (i.e. an anhedonia-like effect) in adolescent females, but not males (Sex × ELS: F[1,63]=6.5, p=0.013). **(E)** In the novelty-suppressed feeding test, ELS produced increased latency to feed (i.e. an anxiety-like effect) in females but not males, (ELS: F[1,63]=4.4, p=0.04; Sex: F[1,63]=11.52, p=0.0012; Sex × ELS: F[1,63]=2.346, p=0.1306). **(F)** In the open field test, ELS did not influence locomotor activity (Sex × ELS: F[1, 63]=0.3390, p=0.5625). In an independent cohort, ELS **(G)** elevated grooming behavior in male but not female adolescent mice in an independent replication cohort (Sex × ELS: F[1, 55]=5.128, p=0.0275), **(H)** increased head-body twitching in adolescent males but not females (Sex × ELS: F[1, 55]=11.30, p=0.0014), and **(I)** increased repetitive pathology Z scores in adolescent males but not females (Sex × ELS: F[1, 55]=16.64, p=0.0001). Two-way ANOVA followed by Sidak’s multiple comparison test: *p<0.05, **p<0.01, ***p<0.001, ****p<0.0001.

The same mice were re-tested in adulthood (**Figure S2**). The effect of ELS on grooming in male mice was lost in adulthood, but social deficits persisted. The depression-like and anxiety-like effects of ELS in females also resolved in adulthood. Locomotor activity was again not affected.

This effect of PND2S ELS on grooming in adolescents is novel, as is the observed sexual dimorphism. We performed an independent replication to confirm the finding. Again, adolescent males but not females showed elevated grooming, replicating the key finding (**Figure 1G**). We also observed increased head-body twitching, which has been interprated as a tic-like manifestation,^38^ in males but not females (**Figure 1H**). To enhance sensitivity of phenotyping repetitive behaviors,^39^ we calculated the repetitive behavioral pathology z-scores, combining grooming and twitching measures. This score was elevated in adolescent PND2S males, but not in females (**Figure 1I**).

Finally, we performed PND2S ELS in a third independent cohort of mice and tested them in adulthood. ELS-induced repetitive grooming, elevation in head-body twitching, and the repetitive behavioral pathology z-scores were not increased by ELS in males or females, confirming the specificity of these effects to adolescents (**Figure S2**).

### ELS influences activities in cortico-striatal and cortico-septal circuits in a sex-specific manner

To investigate the circuit mechanisms mediating behavioral abnormalities in ELS mice, we injected C57BL6 mice of both sexes with retrograde adeno-associated viruses (AAV) expressing mCherry and EGFP into the dorsal striatum (dStr) and lateral septum (LS), respectively, at ∼PND21 (**Figure 2A**). To enhance activity within cortical projections that may underlie ELS-induced social deficits, these mice were either presented an age- and sex-matched stranger mouse in their home cages for 5 minutes or left undisturbed.^40^ Mice were perfused 90 minutes later, and their brains processed for immunohistochemistry. We focused on medial prefrontal cortex (mPFC) and examined co-localization between c-fos, a marker of neural activity, and mCherry (which labels cortico-striatal projection neurons) or EGFP (which labels cortico-septal projection neurons; **Figure 2B**).

**Figure 2.**
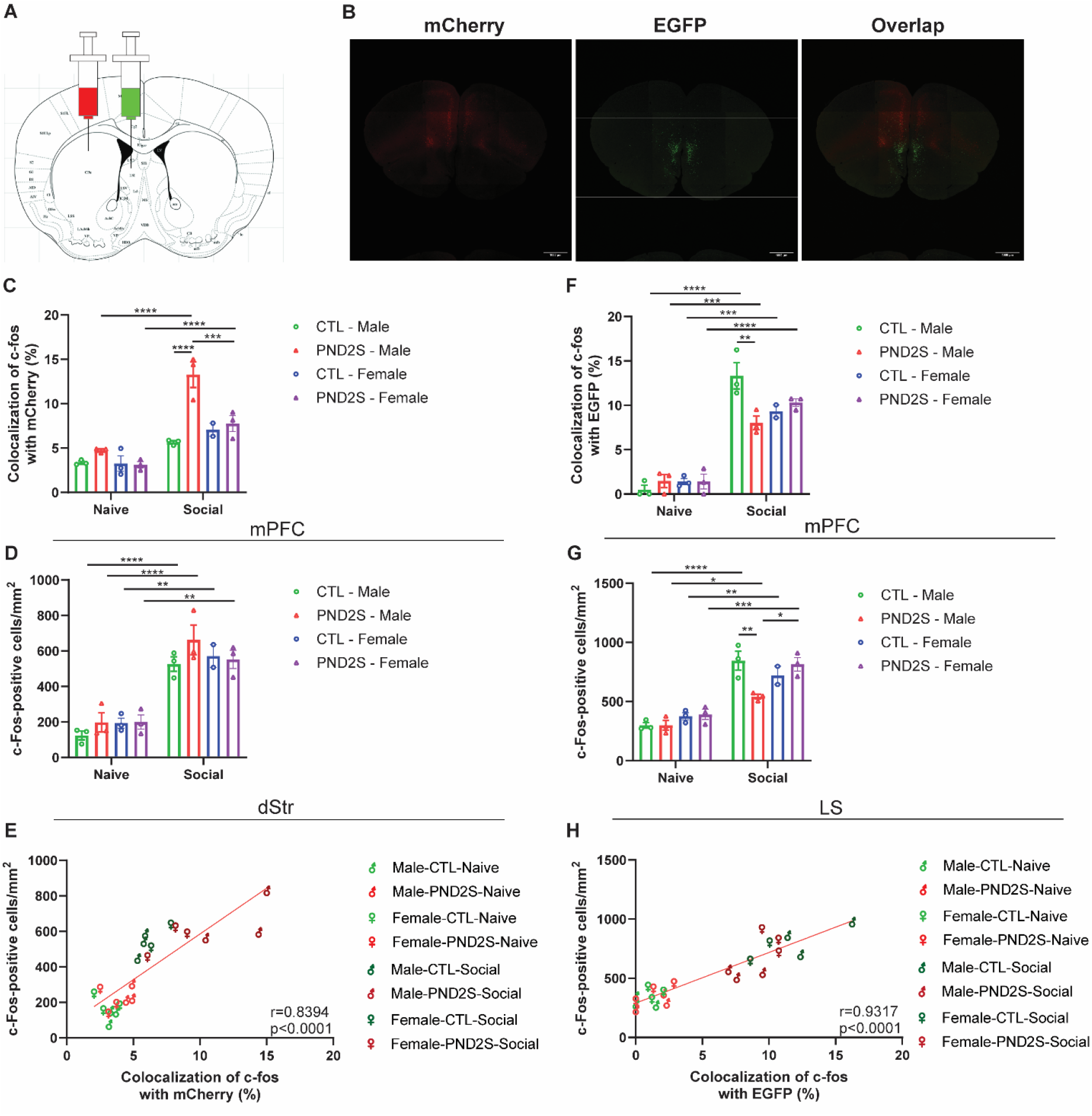
Effects of ELS on cortico-striatal and cortico-septal activity following social stimuli. **(A)** Schematic illustration showing dStr/LS double retrograde tracing strategy. **(B)** Representative images showing cortico-striatal projection neurons (mCherry+) and cortico-septal projection neurons (EGFP+). **(C)** ELS significantly elevated colocalization of c-fos with mCherry-labeled cortical projection cells following 5-minute social encounter with a stranger mouse (Social x Sex x ELS: F[1,15]=6.483, p=0.0224; Sex x ELS: F[1,15]=15.74, p=0.0012; Social x ELS: F[1,15]=11.00, p=0.0047). **(D)** Social stimuli increased # of c-fos+ cells in the dorsal striatum in control and ELS mice of both sexes (Social: F[1,15]=120.5, p<0.0001). **(E)** *c-fos* in cortico-striatal projection neurons was positively correlated with striatal *c-fos* expression in the context of ELS and social stimuli. **(F)** ELS blunted the increase in colocalization of c-fos with EGFP-labeled cortico-septal projection cell induced by social stimuli in males but not females (Social x Sex x ELS: F[1,15]=9.837, p=0.0068; Sex x ELS: F[1,15]=5.275, p=0.0364; Social x ELS: F[1,15]=5.915, p=0.0377). **(G)** ELS blunted the elevation in # of c-fos+ cells in the lateral septum induced by social stimuli in males (Social x Sex x ELS: F[1,15]=7.620, p=0.0146; Sex x ELS: F[1,15]=8.772, p=0.0097). **(H)** *c-fos* in cortico-septal projection neurons was positively correlated with lateral septal *c-fos* expression in the context of ELS and social stimuli. Pearson’s r for E, H. Three-way ANOVA followed by Sidak’s multiple comparison test: *p<0.05, **p<0.01, ***p<0.001, ****p<0.0001.

mCherry+ cells were distributed across the anterior cingulate cortex (ACC), prelimbic cortex (PrL), and infralimbic cortex (IL); EGFP+ were largely restricted to the IL. There was minimal colocalization between mCherry+ and EGFP+ cells (**Figure 2B**), suggesting that distinct populations of cortical neurons project to dStr and LS. ELS significantly elevated social encounter-induced colocalization between c-fos and mCherry (marking cortico-striatal neurons) in males but not females (**Figure 2C**). Social stimuli increased total c-fos+ cells in the striatum, the target of the mCherry+ cortical neurons in both males and females, regardless of ELS (**Figure 2D**). c-fos in cortico-striatal projection neurons was positively correlated with striatal c-fos (**Figure 2E**), suggesting of a functional cortico-striatal circuit.

In contrast, colocalization between c-fos and EGFP (marking cortico-septal projection neurons) was reduced after social stimulation in PND2S ELS male adolescents, but not females (**Figure 2F**). c-fos+ cells in the LS were increased following social stimuli; this increase was similarly attenuated in ELS males but not females (**Figure 2G**). There was a positive correlation between c-fos in cortico-septal projection neurons and c-fos in the LS (**Figure 2H**). Together, these results suggest hyperactivity in the cortico-striatal circuit and hypoactivity in the cortico-septal circuit after PND2S ELS in adolescent males in response to a social stimulus, with no effect of ELS in females.

### Cortico-striatal projection neurons modulate ELS-induced repetitive behaviors and social deficits in males and anhedonia in females

To determine whether cortico-striatal projection neurons regulate repetitive behaviors and social behaviors following ELS, we utilized a retrograde viral tracing strategy and viruses expressing designer receptors exclusively activated by designer drugs (DREADDs) to bidirectionally manipulate the activity of cortico-striatal projection neurons. We injected retrograde virus expressing Cre into the dStr and virus expressing excitatory DREADD hM3D(Gq) or inhibitory DREADD hM3D(Gi) under Cre control, or control virus expressing mCherry, into the mPFC of C57BL/6 mice, with or without PND2S ELS, around the time of weaning (∼PND21). Two weeks after surgery (∼PND35), mice tested in the open field test (OFT) and the three-chamber sociability test (TCST) on two consecutive days, 1 hour after injection of saline or clozapine N-oxide (CNO); repetitive behaviors and sociability were assessed. Another round of OFT and TCST was performed with counterbalanced saline/CNO administration two days later to maintain a within-subjects design (**Figure 3A**).

**Figure 3.**
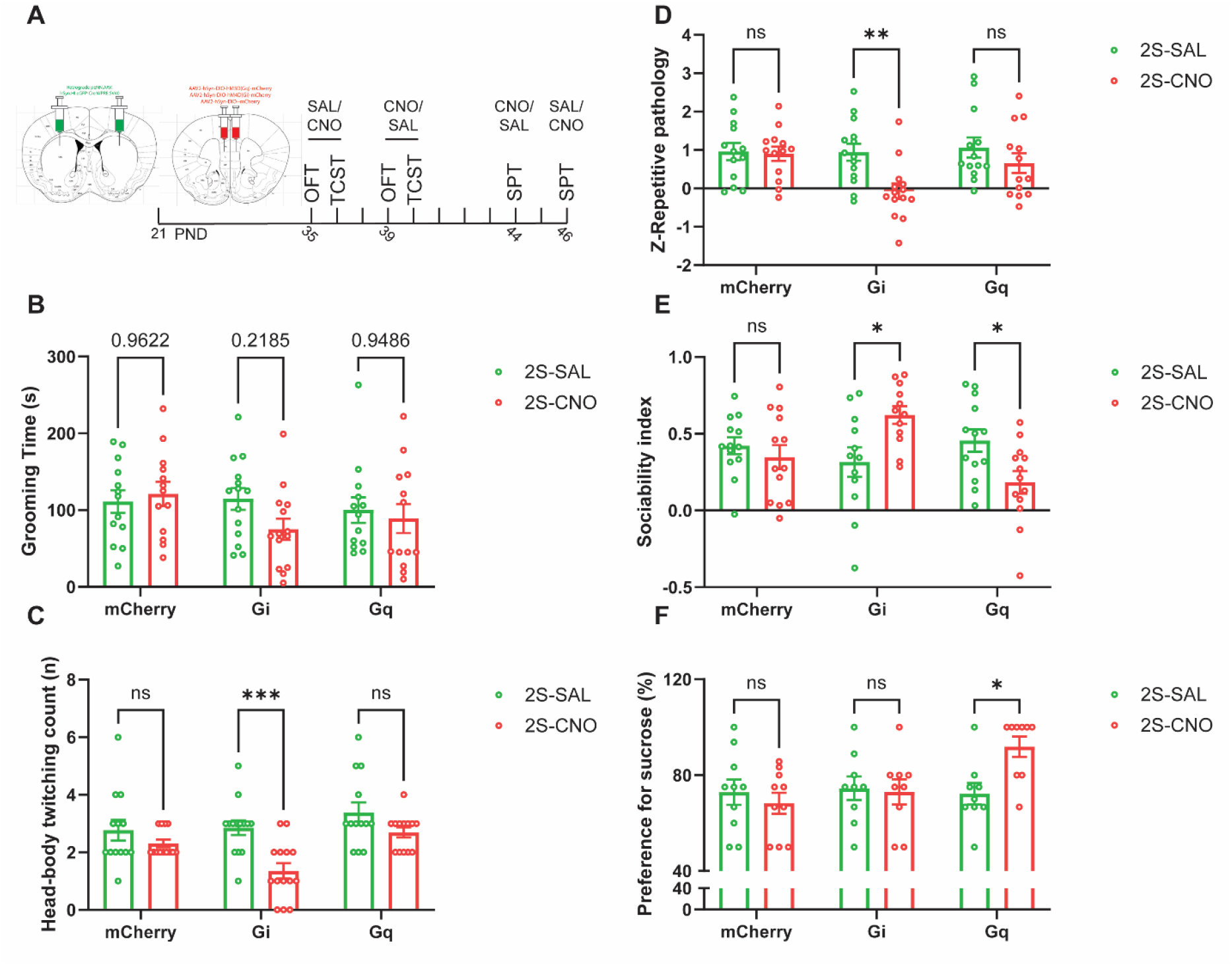
Chemogenetic manipulation of cortico-striatal projection neurons modulate repetitive behavioral pathology, sociability and anhedonia. **(A)** Schematic timeline of chemogenetic manipulation experiments. Chemogenetic inhibition of cortico-striatal projection neurons in adolescent ELS males **(B)** reduced grooming time at trend level (DREADD × CNO: F[2,37]=1.245, p=0.2998), **(C)** significantly reduced head-body twitching (DREADD: F[2,37]=5.680, p=0.0071; CNO: F[1,37]=17.14, p=0.0002), and **(D)** repetitive pathology Z scores (DREADD×CNO: F[2,37]=2.597, p=0.0880; CNO: F[1,37]=7.939, p=0.0077). Chemogenetic manipulations of cortico-striatal projection neurons in adolescent ELS males **(E)** bidirectionally regulated sociability index (DREADD×CNO: F[2,37]=9.908, p=0.0004). **(F)** Chemogenetic activation of cortico-striatal projection neurons rescued anhedonia in adolescent ELS females (DREADD × CNO: F[2,25]=4.254, p=0.0257). Two-way ANOVA followed by Sidak’s multiple comparison test: *p<0.05, **p<0.01, ***p<0.001.

For the OFT, we observed a main effect of PND2S ELS on repetitive grooming, head-body twitching, and repetitive behavioral pathology z-scores in males (**Figure S3A-C**), replicating the effects seen previously (see **Figure 1**). Focusing on the ELS mice, we observed that inhibition of cortico-striatal projection neurons numerically reduced grooming time (**Figure 3B**), significantly reduced head-body twitching (**Figure 3C**), and significantly reduced repetitive behavioral pathology z-scores (**Figure 3D**). Activation of cortico-striatal projection neurons did not affect any of these measures.

In the TCST, we again replicated the effect of PND2S ELS on sociability in adolescent males (**Figure S3D**). Within the ELS group, activation of cortico-striatal projection neurons exacerbated the reduced sociability in males, while inhibition attenuated it (**Figure 3E**). Interestingly, activation and inhibition of cortico-striatal projection neurons did not affect either repetitive behaviors or social behaviors in control mice (**Figure S3A-D**), suggesting that ELS developmentally alters this circuit to produce lasting dysregulation.

Finally, we tested adolescent female mice in the sucrose preference test (SPT) (**Figure 3A**). Mice were treated with either saline or CNO 1 hour before the start of dark phase, during which they were able to freely choose between water or 1% sucrose solution for 2 hours. The same test was performed two days later with counterbalanced CNO or saline treatment. Consistent with our initial observation (**Figure 1**), PND2S ELS produced anhedonia in adolescent females (**Figure S3E**). Within the ELS groups, activation of cortico-striatal projection neurons rescued the anhedonic phenotypes in females (**Figure 3F**); inhibition of these neurons had no effect. Neither activation nor inhibition of these neurons affect sucrose preference in unstressed mice (**Figure S3E**). Together, these data suggest that cortico-striatal circuit modulates ELS-induced behavioral phenotypes in a sex-specific manner.

### Cortico-septal projection neurons modulate ELS-induced social deficits in males

To determine whether cortico-septal projection neurons regulate social behaviors following PND2S ELS, we injected retrograde virus expressing Cre into the LS and virus expressing excitatory DREADD hM3D(Gq) or control virus expressing mCherry into the mPFC of C57BL/6 male mice, with or without ELS, on approximately PND21 (**Figure 4A**). Cortico-septal circuit activation had no effect on repetitive grooming or repetitive pathology z-scores; interestingly, it did reduce twitching, in both ELS and control mice (**Figure 4B-D; Figure S4A-C**). In the TCST, CNO activation of cortico-septal projection neurons rescued the reduced sociability seen in the ELS group (**Figure 4E, Figure S4D**). These results indicate that cortico-septal circuit modulate ELS-induced social deficits and twitching behaviors but not repetitive grooming in male mice.

**Figure 4.**
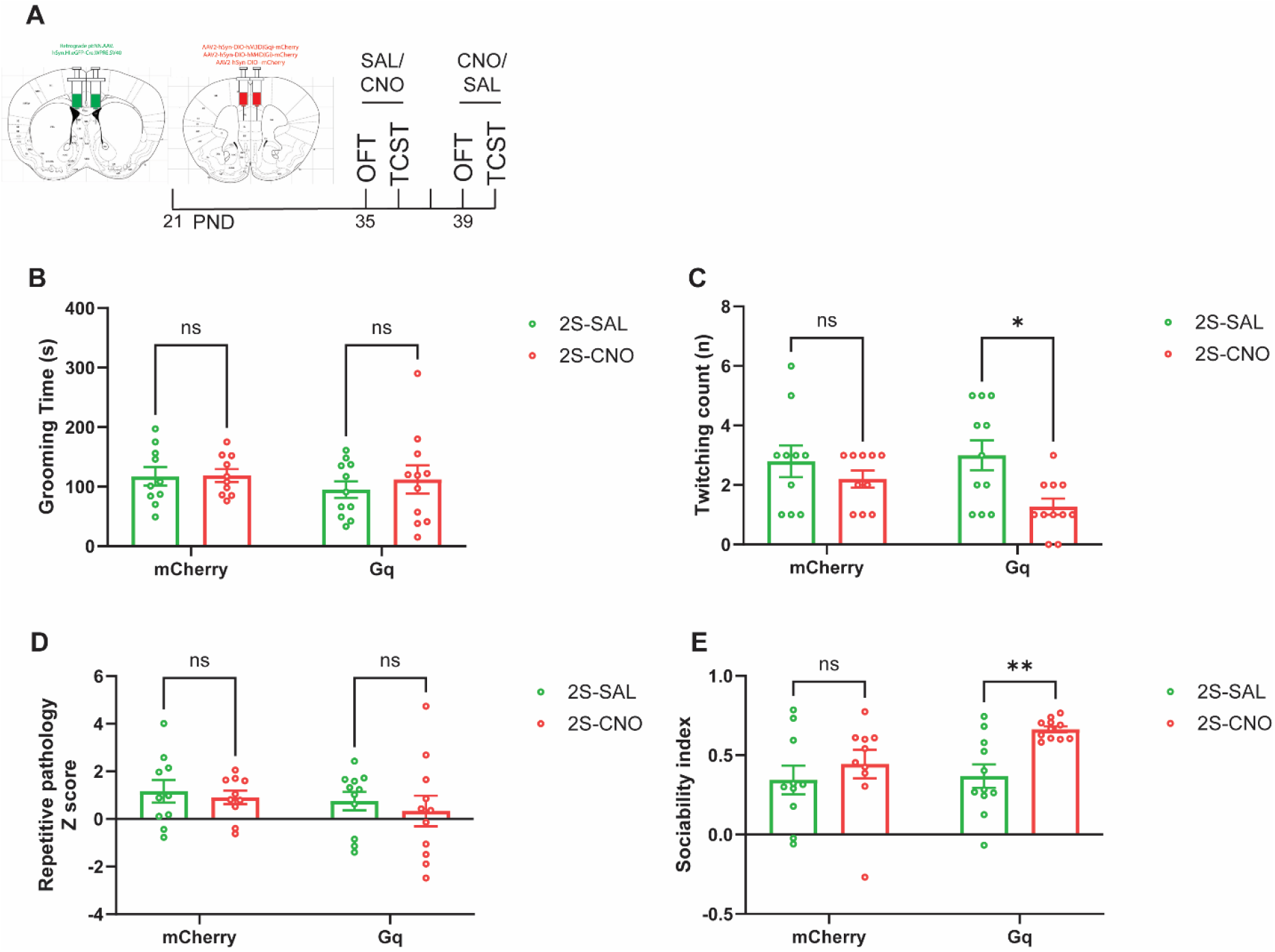
Chemogenetic manipulation of cortico-septal projection neurons modulate sociability. **(A)** Schematic timeline of chemogenetic manipulation experiments. Chemogenetic activation of cortico-septal projection neurons in adolescent ELS males did not influence **(B)** grooming time but reduced **(C)** head-body twitching (CNO: F[1,19]=6.226, p=0.0220) without affecting **(D)** repetitive pathology Z scores. Chemogenetic activation of cortico-septal projection neurons in adolescent ELS males **(E)** rescued deficits in sociability (CNO: F[1,19]=9.219, p=0.0068). Two-way ANOVA followed by Sidak’s multiple comparison test: *p<0.05, **p<0.01.

## DISCUSSION

We demonstrate a novel double dissociation in the effects of ELS, which produces repetitive behavioral pathology and social deficits in male but not female adolescent mice, but depression- and anxiety-like behaviors in females but not males. These effects were replicated in three independent cohorts of mice. While ELS in mice cannot be considered to be a comprehensive model for any neuropsychiatric disorder, this double dissociation is strikingly parallel to epidemiological findings regarding stress-influenced neuropsychiatric conditions. Developmental stress has been associated with ASD,^6-14^ TS,^15-17,41,42^ depression,^43^ and anxiety disorders.^44^ ASD and TS, which are characterized by repetitive behaviors (stereotypies and tics) and often by social deficits, are more common in males,^18,19^ while depression and most anxiety disorders are more common in females.^5^ Furthermore, we find that repetitive behavioral pathology in male mice improves with age, but social deficits persist. Stereotypies improve with age in some individuals with autism,^45,46^ and tics improve in most individuals with TS,^47^ but social deficits in autism tend to persist. Therefore, this ELS paradigm recapitulates key aspects of the phenomenology, sex bias, and developmental trajectory of several neuropsychiatric conditions. We speculate that the underlying mechanisms may prove to be of relevance to the sexually dimorphic pathophysiology of mental illness. Irrespective of its potential connection to human illness, this novel finding illustrates how identical environmental manipulations can have markedly disparate effects in males and females. This paradigm provides an opportunity to clarify mechanisms associated with the sexually dimorphic effects of developmental stress, and how they may contribute to neurodevelopmental and neuropsychiatric pathology.

The mPFC projects to multiple subcortical structures, including the dorsal striatum and the lateral septum. Different circuits may contribute to different aspects of the complex symptom profile of conditions such as ASD and TS,^48-50^ as well as to the development of depression-relevant phenotypes.^51^

Cortico-striatal projections are implicated in the generation and regulation of stereotypies and tics. Individuals with autism show greater activation in the cortex and caudate nucleus during tasks requiring sensorimotor control.^52^ Activity in frontal cortex and caudate is similarly increased during tic suppression; correlated activity suggests that these regions constitute a pathologically dysregulated circuit in individuals with tics/TS.^53^ Similarly, the anterior cingulate cortex is activated prior to tic onset.^54^ Synaptic dysfunction within cortico-striatal circuit has been observed in several genetic animal models of ASD.^55-57^ After ELS, we find markedly increased activity in striatum-projecting cortical neurons after activation of the circuit by a social stimulus, in adolescent ELS males but not in females. Activity in these cortico-striatal projection neurons correlates with activity in the dStr.

Chemogenetic inhibition of cortico-striatal projection neurons ameliorates ELS-induced repetitive behavioral pathology in adolescent males. This parallels a previous study showing that repeated stimulation of orbitofronto-striatal projections leads to development of repetitive grooming in stress-naive mice.^28^ However, we see blunted repetitive behaviors after inhibition of this circuit only in ELS mice; there is no effect in stress-naïve mice. This suggests that the modulation of repetitive behaviors is not a general function of this circuit but manifests only in the context of developmental dysregulation associated with pathology. Interestingly, chemogenetic activation of these neurons does not influence repetitive behaviors, in either ELS or control mice. This is at odds with aforementioned work; this may be attributed to multiple differences between the two studies, including modulation methods (acute chemogenetic modulation vs. repeated optogenetic modulation), cortical region (mPFC vs orbitofrontal cortex), and striatal region (dStr vs. vmStr). Indeed, our previous work show that chemogenetic activation of a tagged subset of mPFC neurons regulated by histamine does not influence grooming behaviors.^29^

Chemogenetic manipulation of cortico-striatal projection neurons also regulates sociability in adolescent ELS males, again without effects in unstressed males. Interestingly, in this case the modulation of behavior is bidirectional: activation of the cortico-striatal projection exacerbates social deficits, while inhibition normalizes sociability. Few previous studies have examined the role of cortical projections to the dorsal striatum in modulating social behaviors.^58^ Indeed, it has long been hypothesized that, in ASD, social deficits are largely regulated by the nucleus accumbens (NAc), perhaps as part of its more general role in reward processing, while repetitive behaviors are mainly regulated by dStr.^59-61^ However, other recent studies have begun to implicate the dStr in social behaviors. Direct manipulations of neurons the dStr influence social functions in addition to repetitive behaviors.^62^ For example, we have found that interneuron depletion in the dStr produces social deficits (interestingly, this effect is again seen only in males).^31^ Activation of D1-expressing spiny projection neurons (SPNs) in the dStr produces social deficits and inhibition of D1-expressing SPNs rescues social deficits induced by prenatal valproate exposure, while activation of D2-expressing SPNs modulates repetitive grooming but not social behaviors.^63^ It is possible that the two categories of behaviors that are regulated by the cortico-dStr projection after ELS (in males) relate to these two categories of SPNs; this remains to be tested.

Interestingly, we found that chemogenetic activation of dStr-projecting cortical neurons rescues anhedonia in adolescent ELS females. This is striking given that we did not see any alteration in the activity of this circuit in ELS females (Figure 1). Neuroimaging studies have revealed structural abnormalities in the dStr (caudate and putamen) of patients with depression.^64-66^ Functional connectivity between cortex and dStr is correlated with depression, and specifically with anhedonia.^67^ The role of the dStr and of the cortico-dStr circuit in depression is not well understood, especially in contrast with the NAc and the cortico-accumbal circuit, which has been extensively studied in this context.^68-70^ However, substantial evidence supports an association between striatal dopamine release and depression and antidepressant treatment. For example, task-related dopamine release in both dStr and NAc is blunted in individuals with anhedonia.^71^ MDD patients show lower dopamine activity in the striatum.^72^ Ketamine, a rapid-acting antidepressant, increases dopamine release in the striatum in both humans and rodents.^73^ At the circuit level, cortical afferents have been found to influence dopamine release in the striatum.^74^ We speculate that the effects of ELS on anhedonia in females may be related to altered regulation of striatal dopamine; further studies are needed to clarify this possibility.

Importantly, the modulatory effects of cortico-striatal circuit manipulations on repetitive behavioral pathology, sociability, and anhedonia are absent in stress-naïve mice. This suggests that ELS developmentally dysregulates the cortico-dStr circuit, leading to lasting abnormalities that may enhance susceptibility to the emergence of dysregulated behavior.

The lateral septum integrates input from the cortex and projects to downstream regions to generate behavioral response to social stimuli. An IL–LS circuit has recently been found to modulate social novelty preference.^75^ We identify a population of LS-projecting cortical neurons that are hypoactive in adolescent ELS males, after circuit activity is enhanced by a social stimulus. Chemogenetic activation of these cortico-septal projection neurons rescues ELS-induced social deficits in adolescent ELS males; yet again, there is no effect in controls. Less is known about this circuitry than about the cortico-dStr circuitry, and mechanistic speculations remain tentative. One recent study suggests that dopamine D3 receptor-expressing neurons in the LS mediate social dysfunction following ELS.^40^ Whether cortical afferents innervate LS D3-expressing neurons to modulate social behaviors remains to be determined.

In summary, we have discovered novel sexually dimorphic effects of ELS on behavioral phenotypes relevant to ASD, TS, and depression. We have identified subcortical regions that receive cortical projections that mediate these effects, with a cortico-striatal circuit essential for repetitive behavioral pathology, social deficits, and anhedonia, and a cortico-septal circuit essential for social deficits. These preclinical findings capture, for the first time, a double dissociation in the effects of developmental stress on neurodevelopment. They set the stage both for further mechanistic work in animals to uncover underlying developmental processes and for clinical studies testing the provocative hypothesis that similar sexually dimorphic effects may contribute to the marked sex differences in the prevalence and, in some cases, the presentation of neurodevelopmental and depressive disorders in humans.

## METHODS

### Mice

Wild-type C57BL/6J mice (Jackson Laboratory) were maintained on a 12 h light/dark cycle with *ad libitum* access to food and water. Animal use and procedures were in accordance with NIH guidelines and approved by the Yale University Animal Care and Use Committees.

### Early life stress

Our ELS paradigm was adapted from that reported in Peña et al.^33^ It combines two stressors, daily maternal separation and chronically reduced nesting material. Pregnant female mice were individually housed 1 – 5 days before giving birth. Litters were pseudo-randomized to receive ELS or standard housing and cage changes, starting on PND2; litters born at the same time were counterbalanced between conditions. ELS pups were removed to a separate cage placed on warming pad, without the mother, for 4 hours daily from PND2 to PND9 (PND2S) or PND10 – PND17 (PND10S). Nesting material was reduced to 1/3 of control in ELS litters, returning to standard housing on PND10 or PND18. Pups were weaned at PND21 into single-sex cages of 2-5 animals.

### Behavioral testing

#### Open field test (OFT)

Mice were placed in an Omnitech monitor apparatus for 30 minutes. Spontaneous exploratory activity was quantified using the breaking of infrared beams. Standard measures include total beam breaks, ambulatory (sequential) beam breaks, stereotypic (repeated) beam breaks, and vertical activity. Mice were videotaped; grooming was scored during the last 10 minutes, blind to experimental condition.

#### Reciprocal social interaction test (RSIT)

An age- and sex-matched stranger mouse of the same strain was introduced to the open field arena at the end of the OFT. Time spent interacting – sniffing (nose-to-nose or nose-to-anogenital), following, pushing past, crawling over or under each other with physical contact, chasing, and so forth – was scored from video over 5 minutes, blind to experimental condition.

#### Three-chamber test for sociability (TCT)

TCT was performed as previously described^76^. This test consisted of four sessions separated by 5 minutes. 1) A mouse was placed in a three-chamber arena with two empty inverted pencil cups in the end chambers and allowed to freely explore for 10 minutes. 2) The mouse was reintroduced to the arena with one clean paper ball placed under each cup and allowed to explore for 10 minutes (novelty). 3) An age-, sex-, and strain-matched unfamiliar mouse (social stimulus) was placed under one cup and an inanimate object (non-social stimulus) under the other. The experimental mouse was placed in the arena and its behavior videotaped for 10 minutes (sociability). Time spent sniffing or engaging with the stimulus (T_S_) and non-social stimulus (T_NS_) was scored over 10 minutes in sessions 3. Social preference indexes, I_SP_ = (T_S_-T_NS_)/(T_S_+T_NS_) was calculated.

#### Sucrose preference test (SPT)

Mice were individually housed and habituated to two bottles of water for 72 hours. One bottle was then filled with 1% sucrose solution and the other with water. The initial weights of the bottles were recorded. The two bottles were weighed again 24 hours later and switched for an additional 24 hours, to control for side bias; they were then weighed again. Sucrose preference was calculated by dividing total sucrose consumption by total liquid consumption (sucrose + water) over 48 hours. A modified SPT over 2 hours was utilized in chemogenetic manipulation experiments. In this version, individually housed mice were habituated to two bottles – one containing 1% sucrose solution and the other water – for 72 hours. Mice were given saline or CNO (2 mg/kg, i.p.) injection 1 hour before dark phase began when they were allowed to freely choose to drink sucrose solution or water for the next 2 hours with the position of bottles switched at 1 hour mark. Sucrose preference was calculated as described above.

#### Novelty suppressed feeding test (NSFT)

Mice were food deprived overnight before testing. Under dim lighting, mice were placed into an open field with a food pellet in the center. Latency to feed was documented.

### Immunohistochemistry for c-fos

To quantify c-fos immunoreactivity after social stimuli, mice were single-housed three days before the testing day. On the testing day, an age- and sex-matched C57BL/6 mouse (social stimulus) was introduced in the cage and allowed to interact with the experimental mouse for 5 min. The naïve mouse was kept in their home cages undisturbed. 90 minutes later, mice were anesthetized using isoflurane and then perfused transcardially with ice-cold 1× PBS (pH 7.4), followed by ice-cold 4% paraformaldehyde in 1× PBS. Brains were postfixed for 48 hours in the same fixative at 4 °C. Coronal sections were prepared on a cryostat (Leica) at 50 μm. For immunohistochemistry, brain sections were incubated in blocking buffer (10% normal donkey serum, 0.3% Triton X-100 in PBS, and 0.02% sodium azide) for 1 hour at room temperature, and then incubated overnight in primary antibodies (chicken anti-GFP, 1:2000, Aves Labs #GFP-1020; rabbit anti-RFP, 1:1000, Rockland #600-401-379; goat anti-c-fos, 1:500, Santa Cruz Biotechnology #SC-52-G) at 4 °C. Sections were then washed in PBS containing 0.3% Triton X-100 for 4 × 10 min and incubated in secondary antibodies (Alexa Fluor® 488 AffiniPure™ Donkey Anti-Chicken IgY (IgG) (H+L), 1:1000, Jackson ImmunoResearch #703-545-155; Donkey anti-Rabbit IgG (H+L) Highly Cross-Adsorbed Secondary Antibody, Alexa Fluor™ 594, 1:1000, ThermoFisher Scientific #A- 21207; Donkey anti-Goat IgG (H+L) Cross-Adsorbed Secondary Antibody, Alexa Fluor™ 647, 1:1000, ThermoFisher Scientific #A-21447) for 1 hour at room temperature. Sections were then washed in PBS containing 0.3% Triton X-100 for 4 × 10 min and then mounted with ProLong Glass Antifad Mountant (P36984, ThermoFisher Scientific). Images were acquired using a confocal microscope (Olympus FluoView FV1000) and analyzed with ImageJ.

### Stereotaxic surgery and viral gene transfer

C57BL6/J mice (∼21 days old) were anaesthetized by intraperitoneal injection with a mixture of ketamine HCl (100 mg/kg) and xylazine (10 mg/kg) and positioned on a stereotaxic instrument (David Kopf Instruments). For dStr/LS double tracing experiment in adolescents, 0.5 μL of retrograde pAAV-Ef1a-mCherry-IRES-Cre (Addgene #55632-AAVrg) and 0.25 μL of retrograde pENN.AAV.hSyn.HI.eGFP-Cre.WPRE.SV40 (Addgene #105540-AAVrg) were infused into the bilateral dStr (from bregma: AP +0.8mm; ML ±1.5mm; DV -2.8mm) and LS (from bregma: AP +0.8mm; ML ±0.4mm; DV -3.0mm), respectively. For cortico-striatal chemogenetic manipulation experiment, 0.5 μL of retrograde pENN.AAV.hSyn.HI.eGFP-Cre.WPRE.SV40 (Addgene #105540-AAVrg) was infused into the bilateral dStr (from bregma: AP +0.8mm; ML ±1.5mm; DV -2.8mm) and 0.3 μL of pAAV-hSyn-DIO-mCherry (Addgene #50459-AAV2) or pAAV-hSyn-DIO-hM4D(Gi)-mCherry (Addgene #44362-AAV2) or pAAV-hSyn-DIO-hM3D(Gq)-mCherry (Addgene #44361-AAV2) was infused into the bilateral mPFC (from bregma: AP +1.8mm; ML ±0.35mm; DV -2.6mm). For cortico-septal chemogenetic manipulation experiment, 0.25 μL of retrograde pENN.AAV.hSyn.HI.eGFP-Cre.WPRE.SV40 (Addgene #105540-AAVrg) was infused into the bilateral LS (from bregma: AP +0.8mm; ML ±0.4mm; DV -3.0 mm) and 0.5 μL of pAAV-hSyn-DIO-mCherry (Addgene #50459-AAV2) or pAAV-hSyn-DIO-hM3D(Gq)-mCherry (Addgene #44361-AAV2) was infused into the bilateral mPFC (from bregma: AP +1.8mm; ML ±0.35mm; DV -2.6mm).

### Chemogenetic manipulation

Clozapine N-oxide (CNO) dihydrochloride (2 mg/kg, Hello Bio #HB6149) or saline was given intraperitoneally 1 hour before OFT, TCST, or SPT.

### Statistical analysis

Statistical analyses were performed using GraphPad Prism (San Diego, California). Normal distribution and equal variances between groups were tested for each experiment. Comparisons for three groups were made using one-way analysis of variance (ANOVA) followed by Dunnett’s multiple comparisons test. Comparisons for four groups or more were made using two-way or three-way ANOVA followed by Sidak’s multiple comparisons test or Tukey’s multiple comparisons test. Correlation was calculated using Pearson’s r. All tests are two-sided. All data are presented as mean ± s.e.m.. Statistical significance is represented as asterisks at *p* values <0.05 (*), <0.01 (**), <0.001 (***), and <0.0001 (****).

## ACKNOWLEDGEMENT

We would like to thank Kyle King and Serena Sim for their technical assistance. This research was supported by the National Institutes of Health (grants NS101104 and MH127259 to CP, Yale/National Institute on Drug Abuse (NIDA) Neuroproteomics Center Pilot Research Project Grant through DA018343 to CJ) and a Tourette Association of America Young Investigator Award (CJ). This work was partially funded by the State of Connecticut, Department of Mental Health and Addiction Services, but this publication does not express the views of the Department of Mental Health and Addiction Services or the State of Connecticut. The views and opinions expressed are those of the authors.

## AUTHOR CONTRIBUTIONS

CJ and CP designed the study. CJ conducted the experiments, acquired, and analyzed the data. IRS and CM analyzed the data. CJ and CP wrote the manuscript.

