## Supplemental Figures for "Circuit mechanisms underlying sexually dimorphic outcomes of early life stress"

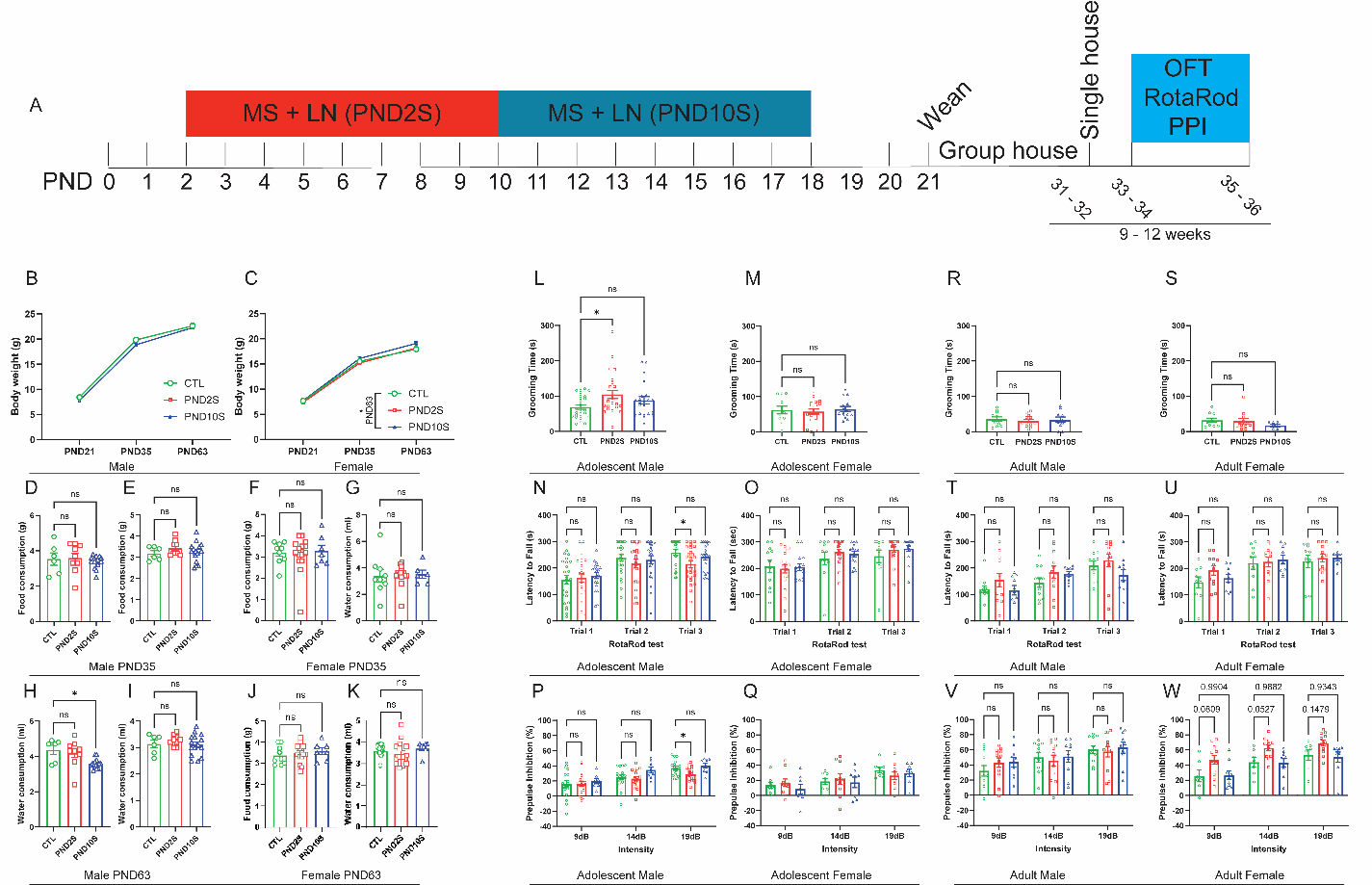

**Figure S1. PND2S produces sexually dimorphic behavioral phenotypes. (A)** Schematic timeline of ELS and behavioral testing. **(B, C)** Influence of ELS on body weight of male and female mice across development. **(D-G)** Influence of ELS on food and water consumptions of male and female mice in adolescence. **(H-K)** Influence of ELS on food and water consumptions of male and female mice in adulthood. PND2S, but not PND10S, elevated grooming time in **(L)** adolescent males (F[2,69]=3.390, p=0.0394) but not **(M)** females (F[2,41]=0.1995, p=0.8199). PND2S, but not PND10S, reduced latency to fall during rotarod test in **(N)** adolescent males (Trial × ELS: F[4,138]=1.572, p=0.1852) but not **(O)** females (F[4,82]=0.4743, p=0.7545). PND2S, but not PND10S impaired prepulse inhibition in **(P)** adolescent males (ELS: F[2,39]=2.059, p=0.1412) but not **(Q)** females (ELS: F[2,23]=0.1518, p=0.8600). ELS did not influence **(R, S)** grooming behaviors, **(T, U)** motor learning skill, and **(V, W)** prepulse inhibition in adult mice of either sex. One-way ANOVA or two-way ANOVA followed by Dunnett's multiple comparisons test: *p<0.05.

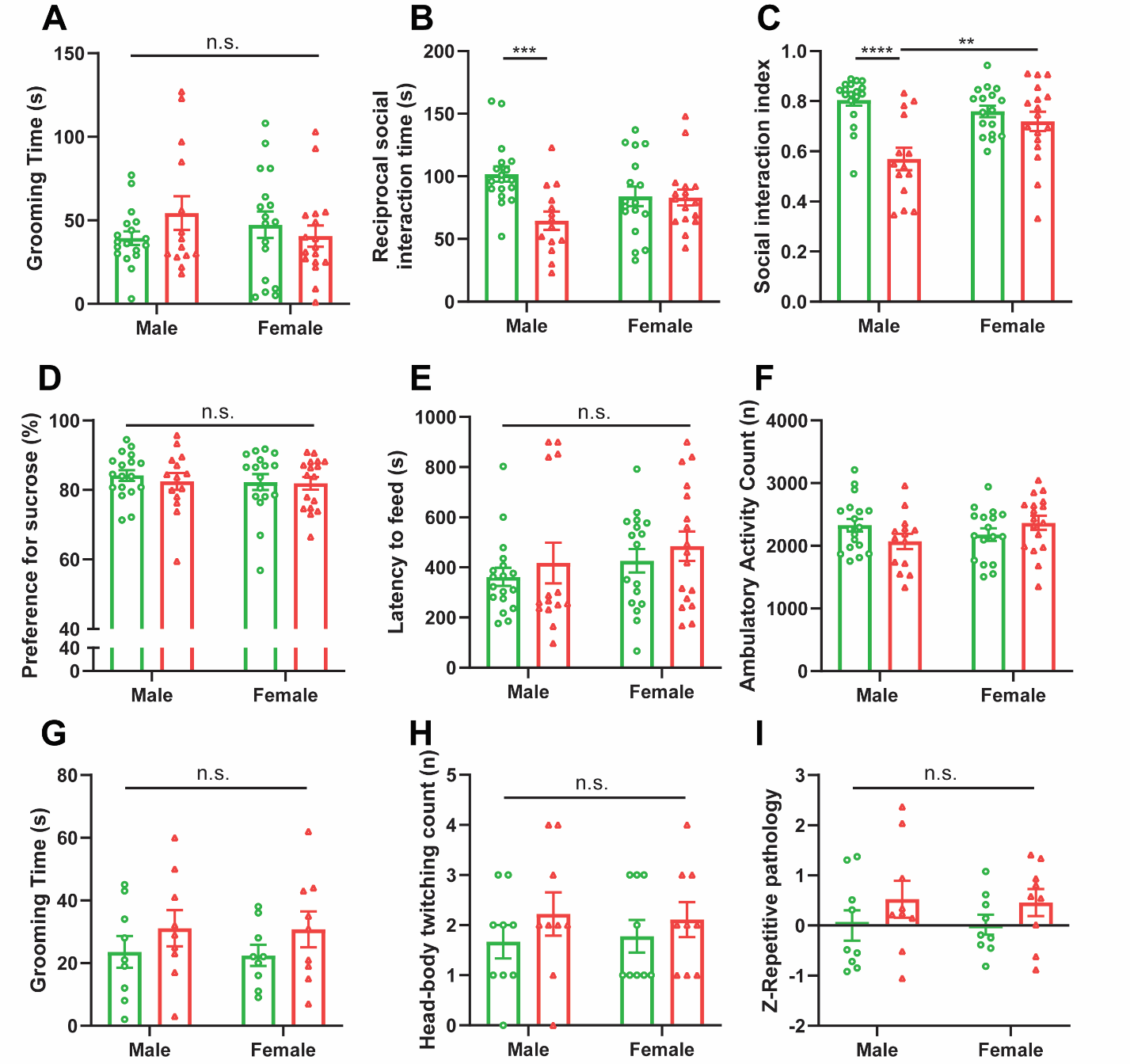

**Figure S2. Behavioral effects of ELS on adult mice. (A)** No effects of sex or of ELS were seen on grooming behaviors in adult mice. **(B)** Reciprocal social interaction was reduced by ELS in adult males but not females (Sex × ELS: F[1, 62]=6.78, p=0.01; ELS: F[1, 62]=7.55, p=0.008). **(C)** In the three-chamber sociability test, ELS reduced social preference in adult males but not females (Sex × ELS: F[1, 62]=9.00, p=0.004; ELS: F[1, 62]=18.0, p<0.0001). **(D)** No effects of sex or of ELS were seen on sucrose preference in adult mice (Sex × ELS: F[1, 62]=0.1105, p=0.7407). **(E)** No effects of sex or of ELS were seen on latency to feed in the novelty-suppressed feeding test in adult mice (Sex × ELS: F[1, 62]=0.0004, p=0.9838). **(F)** A significant interaction between ELS and sex was observed in the locomotor activity (Sex × ELS: F[1, 62]=4.256, p=0.0433). In an independent cohort, ELS did not influence **(G)** grooming behavior in adult mice (Sex × ELS: F[1, 32]=0.006, p=0.9393), **(H)** head-body twitching (Sex × ELS: F[1, 32]=0.0936, p=0.7617), and **(I)** repetitive pathology Z scores (Sex × ELS: F[1, 32]=0.0215, p=0.8842). Two-way ANOVA followed by Sidak’s multiple comparison test: **p<0.01, ***p<0.001, ****p<0.0001.

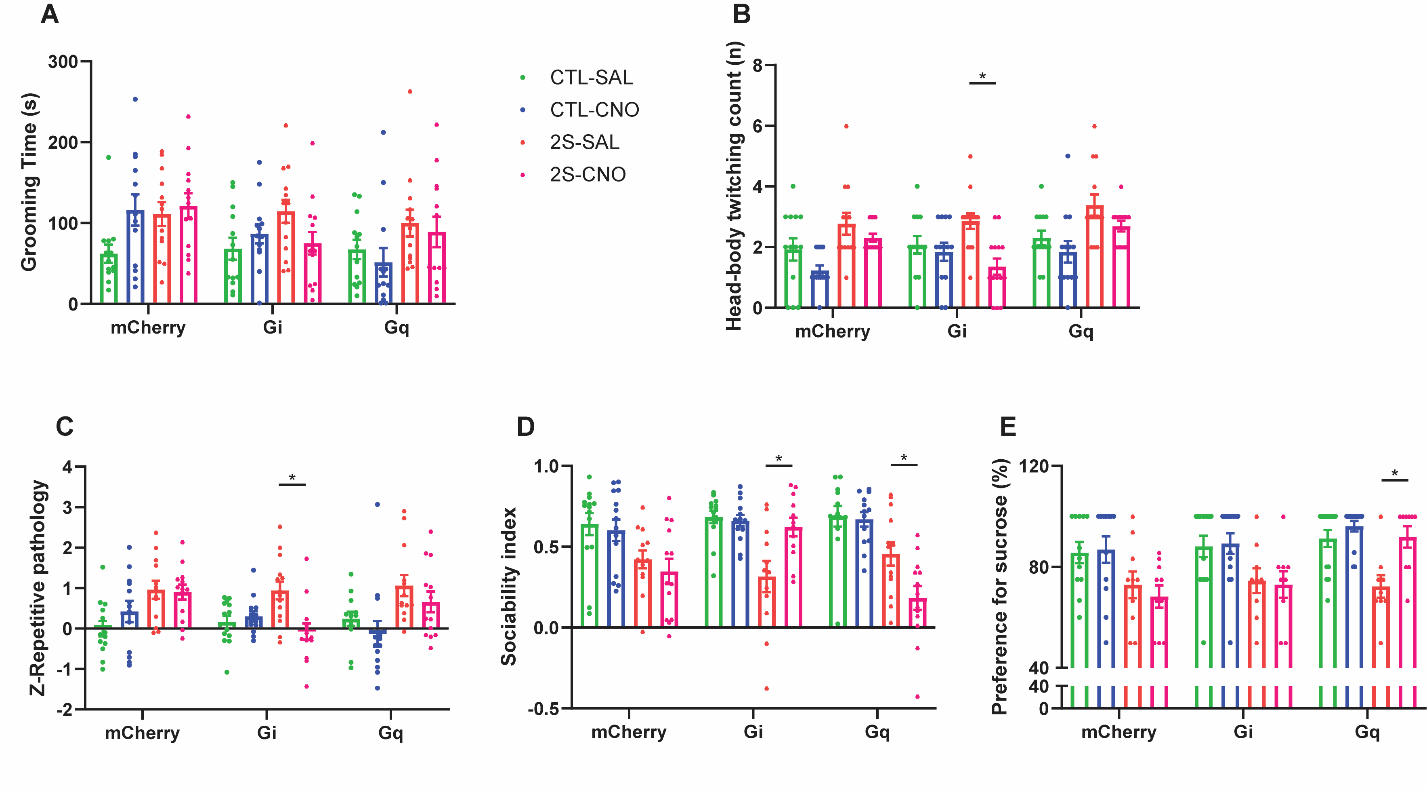
**Figure S3. Chemogenetic manipulation of cortico-striatal projection neurons modulate repetitive behavioral pathology, sociability, and anhedonia.** ELS influenced **(A)** grooming (ELS: F[1,73]=9.886, p=0.0024), **(B)** head-body twitching (ELS: F[1,73]=17.73, p<0.0001), and **(C)** repetitive pathology Z scores (ELS: F[1,73]=13.71, p=0.0004; ELS x CNO: F[1,73]=8.028, p=0.0060) in adolescent male mice that received DREADD infusions. **(D)** Chemogenetic manipulation of cortico-striatal projection neurons bidirectionally regulated sociability (ELS: F[1,73]=47.65, p<0.0001; DREADD x CNO: F[2,73]=5.76, p=0.0047; DREADD x ELS x CNO: F[2,73]=5.739, p=0.0048). **(E)** Chemogenetic activation of cortico-striatal projection neurons rescued anhedonia in adolescent female mice (ELS: F[1,62]=27.61, p<0.0001; DREADD x CNO: F[2,62]=3.525, p=0.0355). Three-way ANOVA followed by Tukey’s multiple comparisons test: *p<0.05.

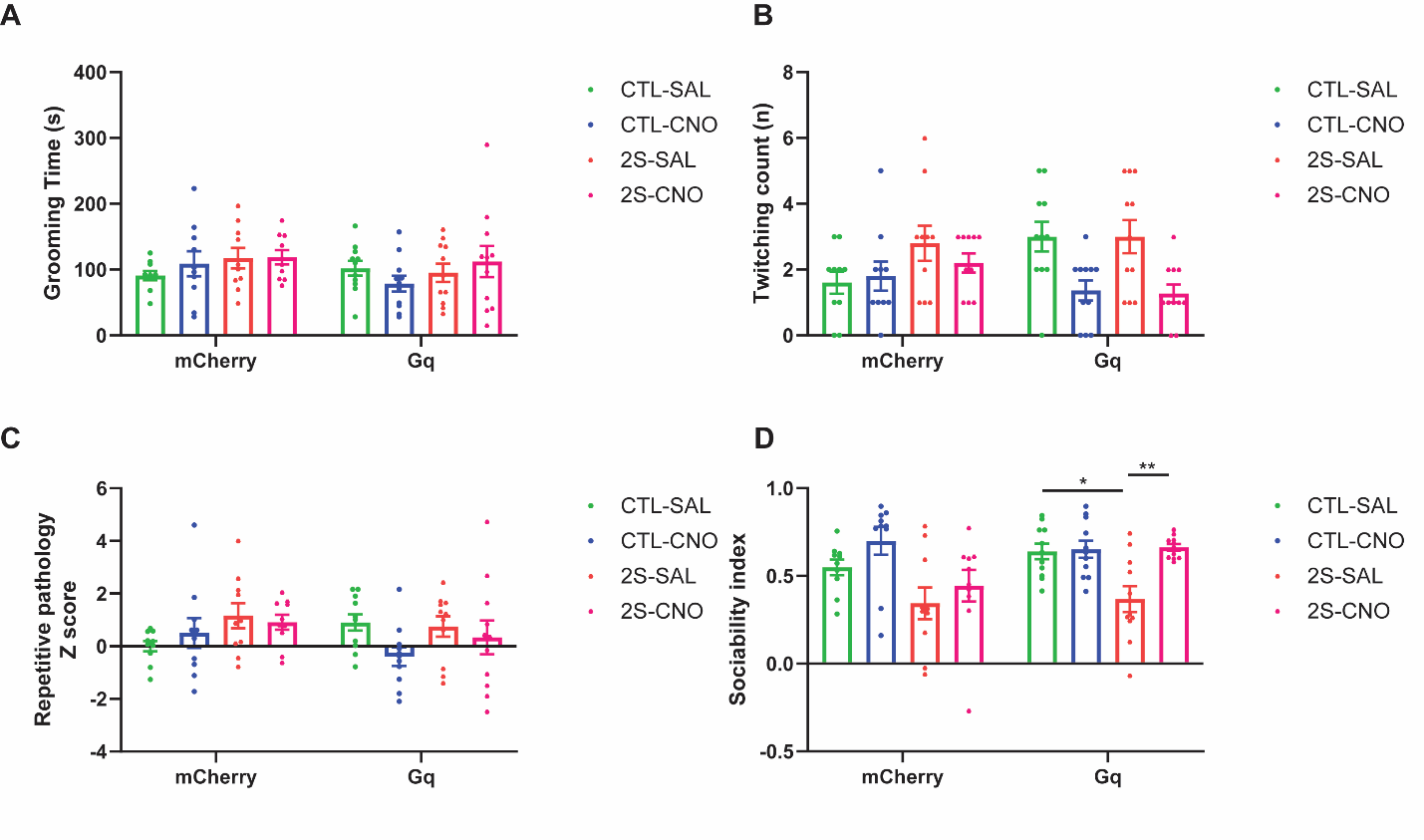

**Figure S4. Chemogenetic manipulation of cortico-septal projection neurons modulate sociability.** Chemogenetic activation of cortico-septal projection neurons in adolescent ELS males did not influence **(A)** grooming time and **(C)** repetitive pathology Z scores, but reduced **(B)** head-body twitching. Chemogenetic activation of cortico-septal projection neurons in adolescent ELS males **(D)** rescued deficits in sociability ELS: F[1,38]=12.75, p=0.0010; DREADD x ELS x CNO: F[1,38]=4.266, p=0.0457). Three-way ANOVA followed by Sidak’s multiple comparison test: *p<0.05, **p<0.01.
